# Sucrose-mediated translational stalling involves a conserved ribosomal pocket

**DOI:** 10.1101/2023.08.27.554957

**Authors:** Sjors van der Horst, Robert Englmeier, Johannes Hanson, Sjef Smeekens, Friedrich Förster

## Abstract

Within eukaryotes, 20-50% of the mRNAs contain short open reading frames (uORFs) located upstream of the main ORF. A significant fraction of these uORFs encode conserved peptides (CPuORFs) that regulate translation in response to specific metabolites. A well-studied example includes uORF2 of the plant growth inhibiting transcription factor bZIP11. Elevated intracellular sucrose levels lead to ribosome stalling at the stop codon of uORF2, thus reducing bZIP11 protein synthesis. Similar examples can be found in bacteria and animals, e.g. on the bacterial *TnaC* and human *CDH1-NPN\** ORFs that both induce stalling at the stop codon when in the presence of tryptophan and the drug-like molecule PF846, respectively.

In this study, we affinity-purified *in vitro* translated sucrose-stalled wheat ribosomes translating bZIP11-uORF2 and determined the ribosomes’ structures using cryo-electron microscopy. This revealed density inside a pocket in the ribosomal exit tunnel of the plant *Triticum aestivum*, that colocalizes with the binding locations of tryptophan and PF846 in *E. coli* and humans, respectively. We suggest this density corresponds to sucrose. Tryptophan and PF846 mode-of-action was previously proposed to inhibit release factor binding or function. Mutation of the uORF2 stop codon shows that its presence is crucial for sucrose-induced stalling, suggesting that the stalling only manifests during termination and not elongation. Moreover, the structural similarities with tryptophan-induced stalled ribosomes near the peptidyl transferase center indicates that an analogous mechanism of inhibition of release factor function is likely. Our findings suggest a conserved mechanistic framework across different organisms, wherein specific molecules interact with the nascent peptide and ribosome to modulate protein synthesis.

## Introduction

Sucrose is central to plant metabolism, both as the end product of photosynthesis and as the main form of transported carbon. Sucrose is also a key signaling molecule in the control of plant growth and energy metabolism. Sucrose signaling controls production of the C and S1 group basic leucine zipper (bZIP) transcription factor proteins that together are key regulators of metabolism and resource allocation in the plant model *Arabidopsis thaliana* (Hanson et al. 2008; Ma et al. 2011; Thalor et al. 2012; Dröge-Laser and Weiste 2018). S1-group *bZIP* mRNAs possess an upstream open reading frame (uORF) in their 5’ untranslated regions (5’UTR) that encodes an evolutionary conserved peptide sequence (CPuORF) functionally conserved in flowering plants (Weltmeier et al. 2009; Wiese et al. 2004; Peviani et al. 2016). Elevated sucrose levels induce ribosome stalling on the stop codon of this CPuORF, resulting in reduced translation of the downstream bZIP ORF (**Figure 1A**) (Rahmani et al. 2009; Hou et al. 2016; Yamashita et al. 2017).

**Figure 1:**
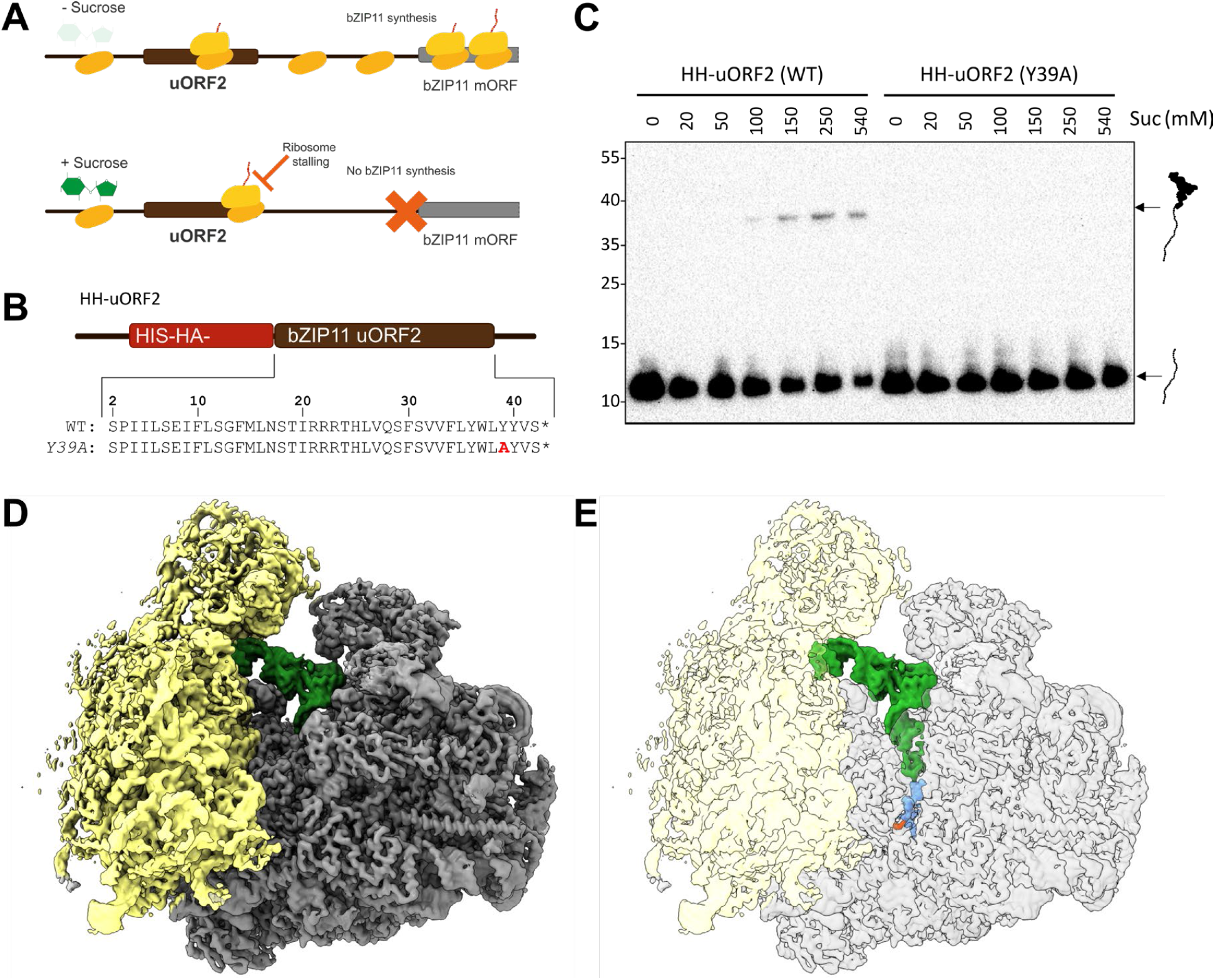
Sucrose induces ribosome stalling on bZIP11-uORF2 in a sequence specific manner. **A**, Schematic representation of sucrose-induced repression of translation of the *bZIP11* mRNA (adapted from (van der Horst et al. 2020)). **B**,**C**, HH-uORF2 (WT or Y39A) mRNAs (**B**) were translated *in vitro* using wheat germ extract for half an hour at 25°C with a varying amount of sucrose. IVT mixtures were separated using SDS-PAGE and analyzed by immunoblot detection using an anti-HA antibody (**C**). Indicated are the full-length peptide (∼12 kDa) and peptidyl-tRNA (∼37 kDa). **D**,**E**, cryo-EM density of sucrose-stalled ribosomes on HHF-uORF2 mRNA, with 40S subunit in yellow, 60S subunit in gray, P-site tRNA in green, uORF2 nascent chain in blue and extra density in orange. In **E**, 40S and 60S subunits are made transparent.

A similar control of translation mechanism by CPuORFs has been observed for other metabolites, including ascorbate, galactinol, phosphocholine and polyamines in plants, arginine in fungi, polyamines in mammals and tryptophan and ornithine in bacteria (van der Horst et al. 2020; Laing et al. 2015; Alatorre-Cobos et al. 2012; Zhu et al. 2018; Uchiyama-Kadokura et al. 2014; Dever et al. 2020). In all these cases a specific amino acid sequence controls translation depending on metabolite concentration. Moreover, many other CPuORFs have been predicted, of which several have shown to induce ribosome stalling (van der Horst et al. 2019; Causier et al. 2022; Hiragori et al. 2022; Takahashi et al. 2020). However, within eukaryotes the molecular sensing mechanism of metabolites remains to be elucidated. Using cryo-EM structural investigations, we show that sucrose binds a conserved binding pocket in the ribosomal exit tunnel and discuss a possible mechanism of the inhibition of translation termination on the S1 group bZIP CPuORFs.

## Results

An *in vitro* translation (IVT) assay with wheat germ extract was used to investigate sucrose-induced ribosome stalling on the CPuORF of S1-group bZIPs. We tested two short constructs containing either a Hexahistidine (H)- and a hemagglutinin (HA)-tag (HH) or a HH-tag followed by a 3x Flag (HHF) tag N-terminally fused to the CPuORF encoding the second uORF (uORF2) of the *Arabidopsis thaliana bZIP11* 5’UTR (**Figure 1B, S1A**). IVT of these mRNAs followed by Western blot analysis using HA antibodies revealed ribosome stalling in the presence of sucrose, as indicated by the peptidyl-tRNA (PtR) band at concentrations as low as 50 mM (**Fig 1C, S1**). As previously reported, for the HHF-uORF2 mRNA a lower PtR band in the absence of sucrose was also observed, but not for the HH-uORF2 mRNA (Yamashita et al. 2017). Sucrose-dependent ribosome stalling is abolished in the uORF2 Y39A (tyrosine to alanine) mutant *in vitro* as previously observed *in planta* (Rahmani et al. 2009; Yamashita et al. 2017) (**Figure 1C, S1**). These results show that sucrose can induce ribosome stalling on uORF2 *in vitro* at physiological concentrations.

To investigate the molecular mechanism behind sucrose-induced ribosome stalling on bZIP11-uORF2, we determined the structure of sucrose-stalled wheat germ ribosome by cryo-EM single particle analysis. To this end, ribosomes translating HHF-uORF2 in the presence of 540 mM sucrose in wheat germ lysate were subjected to affinity purification targeting the N-terminal Histidine affinity tag of the nascent peptide, followed by ultracentrifugation and pellet resuspension (**Figure S2**). Cryo-EM images of the ribosome particles were subjected to several rounds of 3D classification using different masks. In the initial round focusing on the small subunit (SSU), two primary states were identified: 50% of the particles in the unrotated post-translocation state with a P-site tRNA and 17% in a rotated-1 pre-translocation state (**Figure 1D,E, S3 & S4**). The other particles were discarded due to ill-defined rotation state and weak densities for the P-tRNA, and likely represent intermediate states (**Figure S3**). Two subsequent rounds of classification focusing on the tRNAs, resulted in 33% of total particles being classified as the unrotated ribosome and 13% as the rotated ribosome state (**Figure S3**). Several previous studies established that sucrose induces ribosome stalling at the termination step of bZIP11-uORF2 translation (Hou et al. 2016; Yamashita et al. 2017; Yu et al. 2016). Therefore, we focused on the major unrotated ribosome class as the rotated ribosome class cannot have a stop codon in the A-site, as this is occupied by a A/P-tRNA. This unrotated stalled ribosome structure was further refined to a resolution of 3.7 Å. This structure shows a clear P-tRNA density with a continuous density for the nascent peptide, which extends into the exit tunnel, while the A- and E-sites are empty (**Figure 1D-E, 2A, S4**).

Further inspection of the peptide exit tunnel revealed cryo-EM density inside a pocket of the 25S rRNA, which is not explained by the atomic models of the ribosome and is too distant from the nascent chain to be part of it (**Figure 2A-C**). The density connects to the 25S rRNA and the nascent chain. Previous studies on transcript-specific stalling in bacteria and human cell lines induced by a tryptophan (Trp) molecule and the PF846 drug-like molecule, respectively, structurally characterized the binding of the small molecules in the ribosome exit tunnel (van der Stel et al. 2021; Su et al. 2021; Li et al. 2019). The positions of the Trp and PF846 molecules coincide with the position of the additional density identified in the sucrose-stalled ribosome (**Figure 2A**). Closer inspection of the density shows that the overall size and shape is consistent with that of sucrose (**Figure 2B-C**). The density is absent in idle wheat-germ ribosomes in 540 mM sucrose incubated without added mRNA as a control, indicating that this density is associated with translation of uORF2 (**Figure 2B**). Due to the agreement in size of the density of sucrose, its absence in the control dataset and the co-localization with PF846- and Trp inside the stalled ribosomes, we suggest that the density we observe represents sucrose.

**Figure 2:**
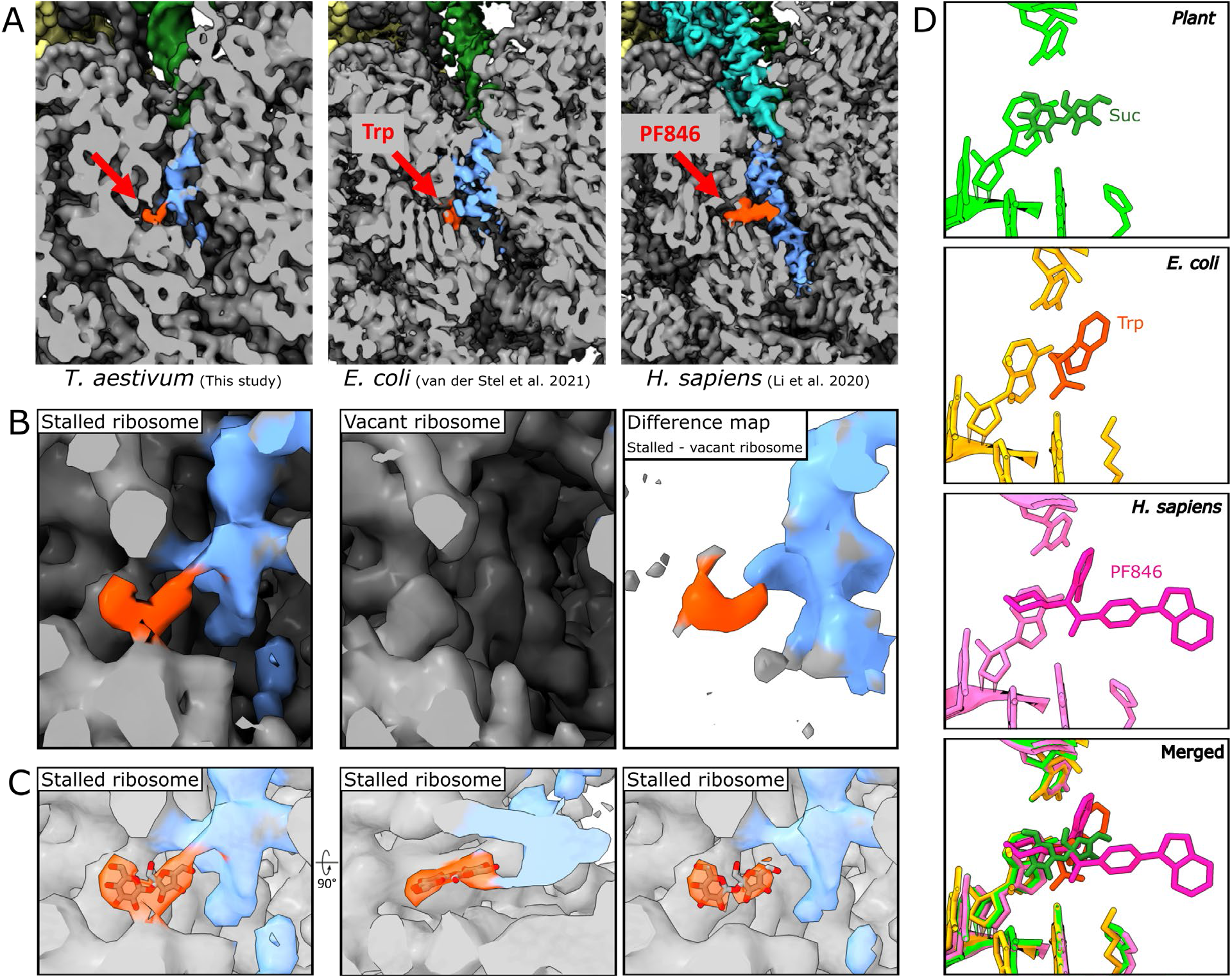
Sucrose binds to a conserved binding pocket in the ribosomal exit tunnel during bZIP11-uORF2 mediated stalling. **A**, Cross-section of cryo-EM maps showing the PTC and the exit tunnel of ribosomes stalled on bZIP11-uORF2 in the presence of sucrose (*T. aestivum*, this study) (left panel), TnaC in the presence of tryptophan (*E. coli*., Van der Stel et al. 2021, EMD-12693) (middle panel) and CDH1-NPN* in the presence of PF846 (*H. sapiens*, Li et al. 2020, EMD-22085) (right panel). Colors as described in **Figure 1**, with the positions of sucrose, tryptophan and PF846 in orange and eRF1 in cyan (right panel). **B**, Map of stalled ribosomes (left panel), and non-translating vacant ribosomes purified from IVT assays where no mRNA was added (middle panel), and difference map of stalled ribosome and vacant ribosome (right panel). **C**, Sucrose molecule fitted into cryo-EM density of sucrose-stalled ribosome, with the contour level threshold increased in the right panel. **D**, Details of sucrose (top), tryptophan (upper-middle), PF846 (lower middle) and all three merged (bottom) with surrounding rRNA nucleotides and His134, Lys90 or His133 of uL22 in tomato, *E. coli* (pdb 7o19) and *H. sapiens* (pdb 6xa1) structures, respectively shows common binding site in molecule induced stalling.

The sucrose-assigned density is located approximately 24 Åfrom the peptidyl transferase center (PTC), which corresponds to approximately 7 (fully stretched) to 16 (alpha-helix) amino acids, while further towards the tunnel exit, the peptide density starts to fade (**Figure 2A**). These 7-16 amino acids of the nascent peptide coincide with the length of the C-terminal stretch that was previously shown to be essential for stalling of the bZIP11-uORF2 peptide (Rahmani et al. 2009; Yamashita et al. 2017). This suggests that peptide-specificity for stalling is likely mediated by multiple side chains that form interactions with the tunnel wall nucleotides and for which we observe multiple connecting densities, a situation similar to the TnaC peptide tunnel wall interaction (**Figure 2A-B**) (Rahmani et al. 2009; Yamashita et al. 2017; van der Stel et al. 2021). The local resolution for the nascent chain is insufficient to unambiguously assign amino acid side chains of bZIP11-uORF2. This prohibits addressing their structural contribution to stalling, but we do observe a connective density between the peptide and sucrose, likely required for stalling (**Figure 2B-C**).

Rigid fitting of the recently published tomato (*Solanum lycopersicum*) ribosome model reveals that the sucrose molecule is in proximity with the conserved rRNA nucleotides C880, G881, A885, G883 and the histidine134 of uL22 (wheat nomenclature) (**Figure 2D**) (Cottilli et al. 2022; Cannone et al. 2002). Sucrose, Trp and PF846 all share negatively charged hydroxy, carboxy or carbonyl groups in proximity to rRNA nucleotide G883, G748, G1584, respectively, indicating the formation of a hydrogen bond, highlighted by the observed density (**Figure 2C-D**). In conclusion, a conserved peptide exit tunnel metabolite binding site appears to be used for molecule recognition by nascent peptides to induce ribosome stalling. In our experiments, the observed density at this position represents a sucrose molecule.

Previous studies showed that sucrose-induced ribosome stalling on bZIP11-uORF2 occurs with the stop codon in the A-site of the ribosome (Hou et al. 2016; Yamashita et al. 2017; Yu et al. 2016). To further investigate the importance of the release factor in sucrose-induced ribosome stalling on bZIP11-uORF2, the stop codon of the HH-uORF2 was changed into an alanine codon and reintroduced into the *bZIP11* 5’UTR 1, 2, 5 or 10 codons downstream (**Figure 3A**). These mRNAs were translated with or without sucrose and analyzed by western blot. When the stop codon was at the wild type position a PtR was visible only in the presence of sucrose and full peptide productions was reduced, indicating ribosome stalling. When the stop codon was positioned further downstream, the stalling was abolished, indicated by both the absence of the PtR when sucrose was present in the IVT reaction and the unrestricted production of the full peptide (**Figure 3A**). This shows that stalling can only occur on bZIP11-uORF2 during termination and not elongation.

**Figure 3:**
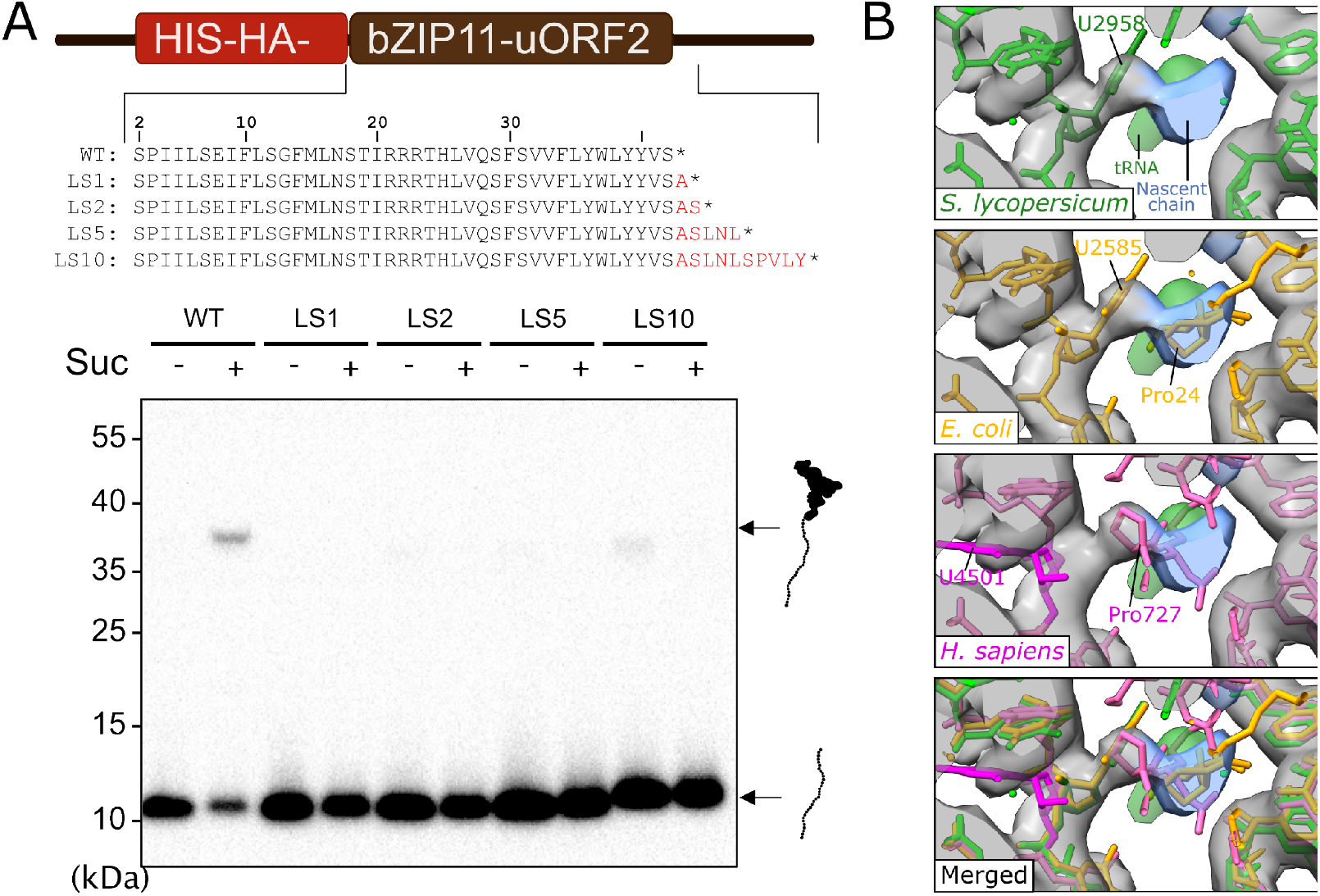
bZIP11-uORF2 mediated stalling relies on the stop codon and the conserved U2956 nucleobase overlaps with that of Trp-stalled U2585. **A**, The stop codon of HH-uORF2(WT) was mutated to an alanine codon and inserted as a late stop codon (LS) one, two, five or ten codons downstream (LS1, LS2, LS5 or LS10, respectively). These RNAs were translated *in vitro* in the presence (+) or absence (-) of 540 mM sucrose and analyzed using SDS-PAGE and immunoblot detection using anti-HA antibody. The peptidyl-tRNA and full peptide are indicated. **B**, Cryo-EM density of the PTC region of the sucrose-stalled ribosome fitted with tomato (green, PDB 7qiz), Trp-stalled *E. coli* ribosomes (orange, PDB 7o19) and PF846-stalled ribosomes on CDH1-NPN* (pink, PDB 6xa1).

Trp-, Ornithine, and PF846-stalled ribosomes all contain the stop codon in the A-site when stalled on the *TnaC, SpeFL* and *CDH1-NPN\** ORFs, respectively (Li et al. 2020; van der Stel et al. 2021; Su et al. 2021; Herrero del Valle et al. 2020). In all cases an uracil nucleotide of the 23S/28S rRNA was suggested to play a role in the inhibition of release factor activity. Interestingly, this uracil, that forms part of the PTC, is conserved between the *E. coli* (U2585), human (U4501), *T. aestivum* (U2956) and *S. lycopersicum* (U2958) ribosomes (Youngman et al. 2004). For the Trp-stalled ribosome it was hypothesized that Pro24 of the TnaC nascent peptide slightly pushes U2585, preventing release factor 2 (RF2) action (van der Stel et al. 2021). By contrast, for the PF846-stalled ribosome U4501 of humans is flipped 90 degrees, with Pro727 occupying the non-flipped position, allowing for eRF1 binding but impaired peptidyl-tRNA hydrolysis (Li et al. 2020). In our structure, the uracil position matches with that of the Trp-stalled ribosome, as well as with that of the translation competent tomato ribosome structure (**Figure 3B**). The sucrose-stalled bZIP11-uORF2 nascent chain overlaps with Pro24 of TnaC and is in close proximity to the U2956 uracil (**Figure 3B**). Additionally, our data did not reveal a ribosome class with bound eRF1 (**Figure S3**). This is also true for the majority of the Trp-stalled ribosomes, albeit that a structure with bound RF2 was observed in a fraction of the particles indicating that RF2 binding was still possible (van der Stel et al. 2021; Su et al. 2021). In conclusion, our results suggest that sucrose-induced stalling on bZIP11-uORF2 may be mediated through inhibition of release factor action at the PTC, similar to Trp- and ornithine-induced ribosome stalling.

## Conclusion

Sucrose is a key energy source, carbon transporter, and signaling molecule in plants. However, the mechanism by which sucrose is sensed to regulate metabolism has remained a mystery for decades. Our cryo-EM study indicates a binding pocket for sucrose inside the ribosome exit tunnel, where sucrose interacts with both the ribosomal RNA and the bZIP11-uORF2 nascent peptide. Mutational analysis of the bZIP11-uORF2 stop codon shows that sucrose-induced ribosome stalling only occurs during translation termination. Additionally, in the sucrose-stalled ribosome we observe structural similarities near the PTC with Trp-stalled ribosomes on *TnaC* ORF, indicating inhibition of eRF1 action. Future investigation including improved local resolution of the nascent chain is necessary to unravel the exact mechanism of the termination stalling.

The identified sucrose binding site overlaps with the binding site of Trp- and PF846 in molecule-induced ribosome stalling during *TnaC* and *CDH1* translation in *E. coli* and humans, respectively. This indicates an ancient functionally conserved small molecule recognition site within the ribosomal exit tunnel that conditionally tunes gene expression at the level of protein production. The 3 billion year evolutionary gap between pro- and eukaryotes and the general abundance of CPuORFs, particularly in plants, suggests that other metabolites may exploit the same binding pocket for translational control.

## Material & Methods

### RNA design and synthesis

DNA templates for *in vitro* transcription were created using Phusion PCR from template plasmids or purchased from GeneArt Strings and directly used for *in vitro* transcription. For plasmid production, SP6::HHF-DP75-uORF2_(WT) construct was ordered from GeneArt Strings (**Supplemental file 1**) and cloned into pJET1.2 plasmid using the CloneJET PCR Cloning Kit (Thermo Fisher Scientific, K1231). To remove the DP75 and 3xFLAG-DP75 regions from the uORF constructs, site-directed mutagenesis was performed using uORF2_FW (TCTCCAATAATACTCAGTGAGATC) as forward primer and 3xFLAG_RV (CTTGTCATCATCGTCCTTGTAG) or GGGGS-HS_RV (GGAACCGCCACCGCCAG) as reverse primer to create SP6::HHF-uORF2 or SP6::HH-uORF2, respectively. DNA templates for *in vitro* transcription were produced by PCR using Phusion Polymerase with the plasmids and SP6_HH_FW (GTTTTTCAGCAAGATTGC) as forward primer and bZIP_FL_RV (ACTGGAGATAAGTTCAGAGATC), or bZIP_Y39A_FL_RV (ACTGGAGATAAGTTCAGAGATCATGAGACATAtgcAAGCCAG) as reverse primer. For the experiment with late stop codons (**Figure 3A**) the DNA templates were purchased (GeneArt™ Custom DNA Fragments, ThermoFisher Scientific) and directly used for *in vitro* transcription (**Supplemental file 1**).

DNA templates were used to produce capped transcripts by *in vitro* transcription using the mMESSAGE mMACHINE™ SP6 Transcription Kit (Thermo Fisher Scientific, AM1340) and the RNA was purified using the MEGAclear™ Transcription Clean-Up Kit (Thermo Fisher Scientific, AM1908) or the GeneJET RNA Cleanup and Concentration Micro Kit (Thermo Fisher Scientific, K0841) for smaller quantities of RNA, all according to the manufacturer’s instructions.

#### *In vitro* translation assay and ribosome purification

For western blot analyses, 20 μl of *in vitro* translation (IVT) reaction was performed in varying concentrations of sucrose using 10 μl wheat germ extract (Promega, L4380), 1.6 μl complete amino acid mixture (Promega, L4461), 0.4 μl Ribolock nuclease inhibitor (ThermoFisher Scientific, EO0382), 6 μl water or concentrated sucrose and 2 μl 125 nM RNA.

For cryo-EM analysis of the vacant ribosome control dataset, 200 μl IVT reactions were briefly spun (5 min at 15,000 *g*) and loaded onto a 1 M sucrose cushion (50 mM Tris-HCl (pH 7.0), 250 mM Potassium acetate (KOAc),25 mM Magnesium acetate (Mg(OAc)2), 2 mM 1,4-dithiothreitol (DTT) and 1M sucrose) followed by ultracentrifugation at 91,000 RPM for 45 minutes in a precooled Beckman Optima TLX with TLA-100.2 rotor. The pellet was resuspended in 50 μl resuspension buffer (20 mM Tris-HCl (pH 7.0), 50 mM KOAc, 10 mM Mg(OAc)2, 1 mM DTT and 150 mM sucrose) and used for cryo-EM analysis.

For affinity purification of stalled ribosomes on HHF-uORF2 mRNA in the presence of sucrose, the IVT was scaled up to 2 x 500 μl reaction in the presence of 540 mM sucrose. Next, the IVT reaction was ultracentrifuged over a 1 M sucrose cushion as described above. The pellets were resuspended in 250 μl ribosome-nascent chain complex (RNC) buffer (50 mM Tris-HCl (pH 7.0), 250 mM KOAc, 25 mM Mg(OAc) 2, 5 mM 2-mercaptoethanol and 540 mM sucrose), combined and incubated with 60 μl cobalt-based magnetic beads (ThermoFisher Scientific, 10103D) for 10 minutes at 4°C, followed by thorough washing with RNC buffer and RNC buffer supplemented with 0.5% Triton-X100. Stalled ribosomes were eluted using RNC buffer with 200 mM imidazole and the eluate was pelleted again through a 1M sucrose cushion. The pelleted ribosomes were resuspended in 30 μl resuspension buffer and stored on ice until they were loaded onto the cryo-EM grid.

### Western Blot analysis

For detection of peptide and peptidyl-tRNA presence, *in vitro* translation reaction or purification intermediates were analyzed using western blots as follows. SDS-PAGE loading buffer (premixed 4 μl Laemmli sample buffer (Bio-Rad) and 1.6 μl Bolt reducing agent (Thermo Fisher Scientific)) was added to the ribosome samples followed by separation on 8% (w/v) Bolt Bis-Tris gel in MES buffer (Thermo Fisher Scientific), followed by transfer onto a 0.45 μm PVDF Immobilon-P membrane (Bio-Chapter 691Rad). The blots were probed with Anti-HA-peroxidase, high affinity antibody (Roche) and chemiluminescence was visualized using a ChemiDoc gel system (Bio-Rad).

### Cryo-EM grid preparation

Cryo-EM grids were prepared by applying 3 μl of ribosome sample at a concentration of 2-4mg/ ml to freshly glow discharged Quantifoil Cu200 R1.2/ 1.3 (HHF-bZIP11-uORF2 + Sucrose) or R2/1 (vacant ribosome control datasets) holey carbon. The grids were flash-frozen using a Vitrobot Mark IV (Thermo Fisher Scientific) with 595 blotting paper (Ted Pella) set to 4 °C, 100% humidity and a blot force of 0 for 4 s, and plunged into a liquid ethane/propane mixture.

### Cryo-electron microscopy data acquisition and processing

The dataset of the sucrose-stalled, HHF-bZIP11-uORF2 translating wheat germ ribosomes was acquired on a 200 kV Talos Arctica (Thermo Fisher Scientific) equipped with a post-column energy filter and a K2 Summit direct-electron detector (Gatan) at Utrecht University. Movies containing 70 frames were acquired with SerialEM 32 at varying defocus of 0.8 - 1.8 μm, at a pixel size of 1.015 Å/ pixel, a dose rate of 4 e-/ pix/ sec and a total dose of 56 e-/ Å. Control datasets of vacant ribosomes were acquired on a 300 kV Titan Krios (Thermo Fischer Scientific) equipped with a post-column energy filter and K3 at the Netherlands Centre for Electron Nanoscopy (NeCEN). Movies containing 42 frames were acquired with SerialEM 32 at varying defocus of 0.8 - 2.2 μm, a pixel size of 0.86 Å/ pixel, a dose rate of 15 e-/ pix/ sec and a total dose of 54 e-/ Å2. Datasets were processed using Relion 3.1 (Scheres 2012). Micrographs were motion-corrected using MotionCor2 (Zheng et al. 2017) and CTF-corrected using gCTF (Zhang 2016). Ribosomes were picked using Relion’s Laplacian-of-Gaussian auto-picking, and false-positive particles were removed by 2 rounds of 2D classification and one round of 3D classification. A high-resolution map of all ribosomes was obtained by alignment using Refine3D. For the separation of the different t-RNA states, masks encompassing the A-, P- and E-tRNA site or just the P-tRNA site were employed using Relion’s background subtraction followed by 3D classification. High-resolution maps of the three translating states were obtained using Relion’s Refine3D, followed by one round of CTF refinement, Bayesian polishing and a final alignment using Refine3D. The maps were sharpened using Phenix’s auto-sharpen (Liebschner et al. 2019).

The tomato (*Solanum lycopersicum)* ribosome model (7qiz, (Cottilli et al. 2022)) was aligned to the sucrose-stalled ribosome using a rigid body fit employing the “fit-in-map” feature in ChimeraX (Pettersen et al. 2021). Using ChimeraX’s volume subtract command with the “minRms true” setting, difference maps were generated between the sucrose-stalled ribosome and the “no mRNA” control ribosome.

## Acknowledgements

We would like to thank Chris Schneijdenberg and Stuart Howes for their support of the Utrecht University CryoEM facility, and Mihajlo Vanevic for his support of the computational infrastructure. We thank NeCEN in Leiden for their support in acquiring the 300 kV dataset. In addition, we thank Julia Bailey-Serres from University of California, Riverside and Sander Roet from Utrecht University for helpful discussions and critical comments on the manuscript. This research was funded by the Dutch Research Council (NWO) (grant no. Vici 724.016.001 to F.F. and ALWOP.2015.115 to S.S.), the Netherlands Electron Microscopy Infrastructure, project number 184.034.014 of the National Roadmap for Large-Scale Research Infrastructure of the Dutch Research Council (NWO) and the European Molecular Biology organization (EMBO grant no. ALTF 264-2021 to S.v.d.H.).

## Author Contributions

R.E., S.v.d.H., F.F., J.H. and S.S. designed experiments. R.E. and S.v.d.H. conducted experiments and analysed the data. S.v.d.H conducted *in vitro* translation assays and affinity purification. R.E. prepared cryoEM grids, acquired the cryo-EM data, performed the single-particle analysis and build the atomic models. R.E. and S.v.d.H. wrote the initial draft of the manuscript, which was edited by F.F., J.H. and S.S.

---

**Figure S1:**
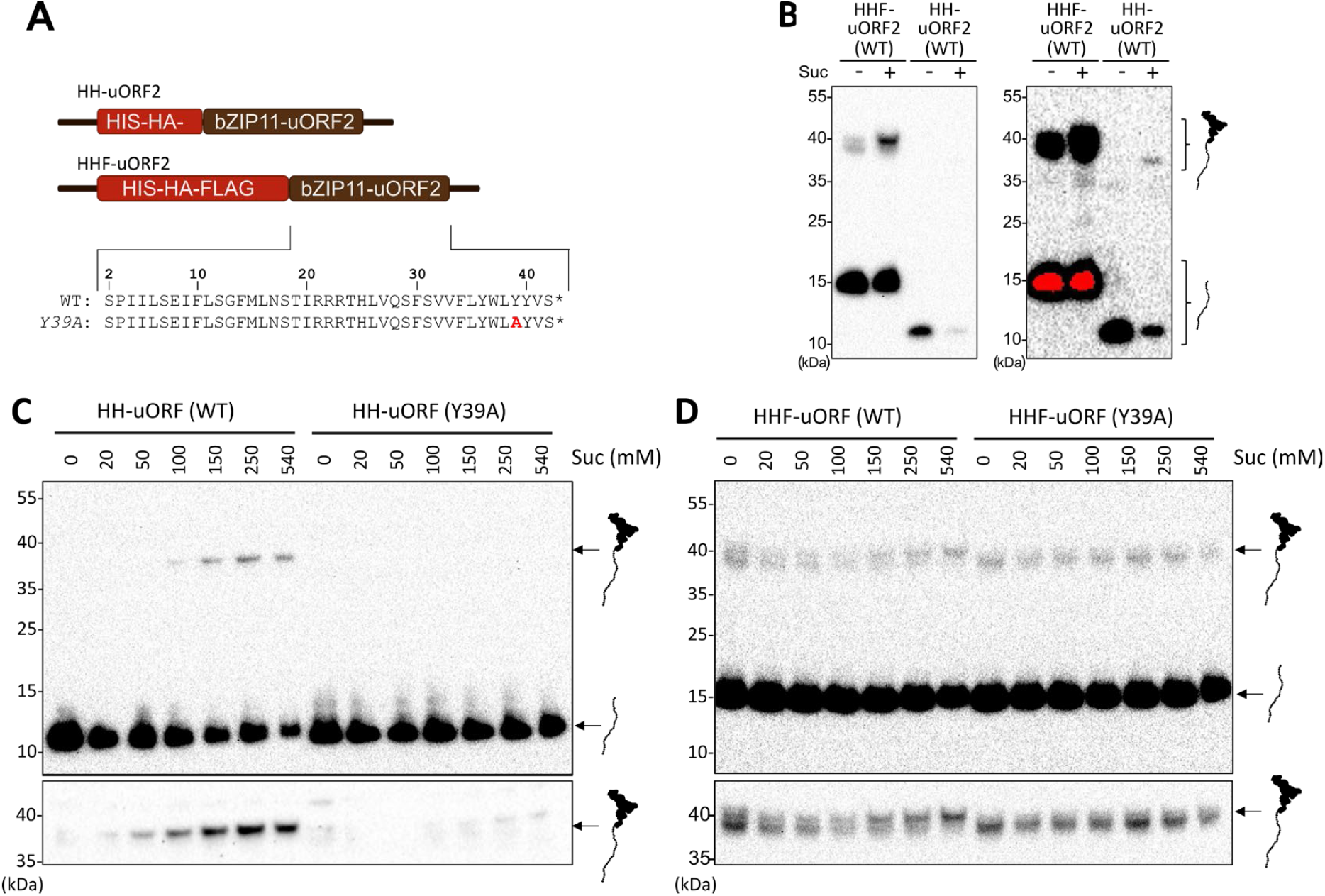
uORF2-bZIP11 induces stalling of ribosomes in a sucrose dependent manner. **A**) The constructs that were used for the *in vitro* translation (IVT) reactions contain different sequences upstream of uORF2: a Hexa-Histidine (HH), Hexa-Histidine-HA-3xFlag (HHF) or Hexa-Histidine-HA-3xFlag followed by a 75 amino acid linker (HHF-DP75)(Yamashita et al. 2017). Wild type and mutant uORF amino acid sequences are shown. **B**) Western blot of IVT reactions with ribosomes translating either the HHF-or the HH-uORF2 transcript, in otherwise identical conditions, in the absence (-) or presence of 540 mM sucrose (+). Both the amount of free peptide (bottom) and the peptidyl tRNA (top) is higher for the HHF transcript. **C**) and **D**) Western blot of IVT reactions with the HH- (**C**) and HHF-uORF2 transcript (**D**) at increasing sucrose concentrations (0 to 540 mM) showing translational stalling for the HH-uORF2 transcript, and a shift from the sucrose-independent peptidyl tRNA to a higher-molecular weight with increasing sucrose concentrations for the HHF-uORF2 transcript. Mutation of tyrosine 39 (Y39) to alanine abolishes stalling. Bottom blots of **C** and **D** and right blot of **B** show the PtR region of the top panel, but with a longer exposure time. Upper blot in **C** and **Figure 1C** are the same images but shown here again for comparison with the HHF-uORF2 transcript.

**Figure S2:**
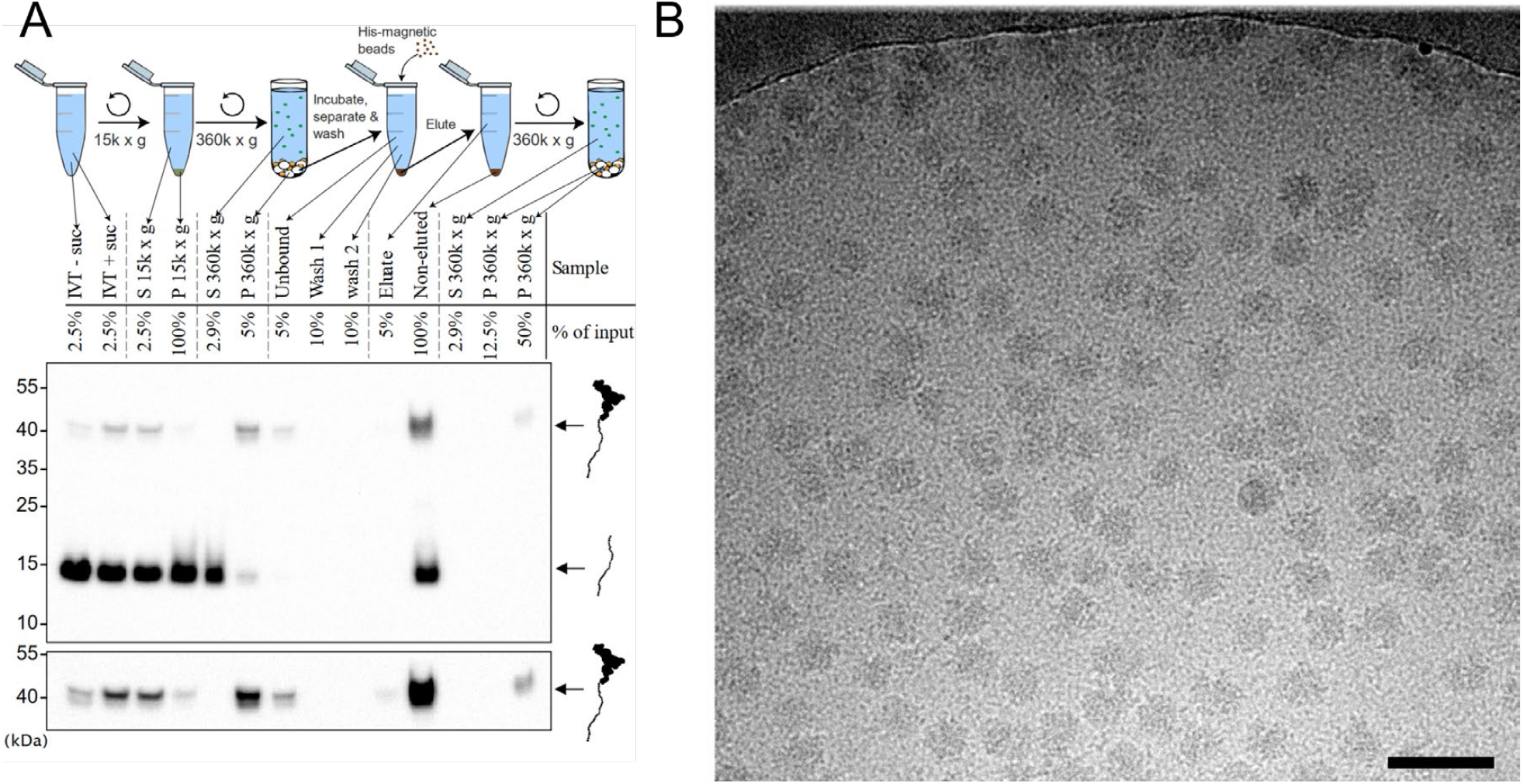
Enrichment of stalled ribosome-nascent chain complexes by affinity purification. **A)** HHF-bZIP11-uORF2 transcripts were translated *in vitro* in the presence of 540 mM sucrose and ribosome-nascent chain complexes (RNCs) were purified by ultracentrifugation and affinity purification using His-magnetic beads. Purification steps were monitored using western blot with anti-HA antibodies. Bottom panel shows the PtR region of the top panel, but with a longer exposure time. Positions of the peptide and PtR bands are indicated. S = Supernatant, P = Pellet. **B)** Cryo EM micrograph of affinity purified RNCs. Scalebar 50 nm.

**Figure S3:**
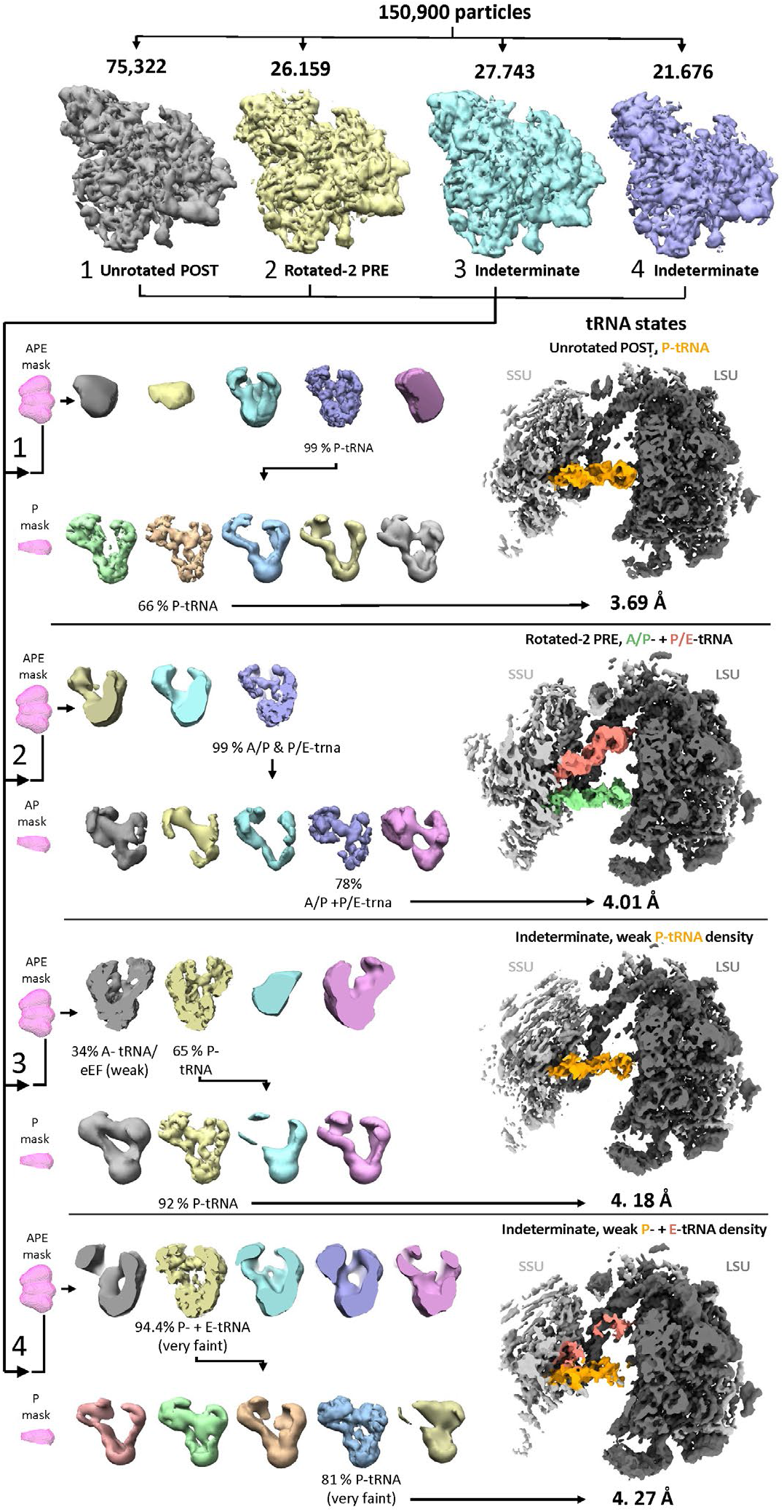
Classification of affinity purified wheat germ ribosomes translating HHF-uORF2 in the presence of sucrose. After initial 2D classification, 3D classification, and 3D refinement, 150,900 ribosome were obtained and subjected to three rounds of classification using masks focusing on the SSU (top, only SSU area is shown), the tRNA binding site (encompassing the A-,P- and E-tRNA site) and the P-site (classes 1, 3,4) or A/P-site t-RNA (class 2). Maps of the tRNA binding site mask in pink are viewed from top, while the respective classes are show from the side.

**Figure S4:**
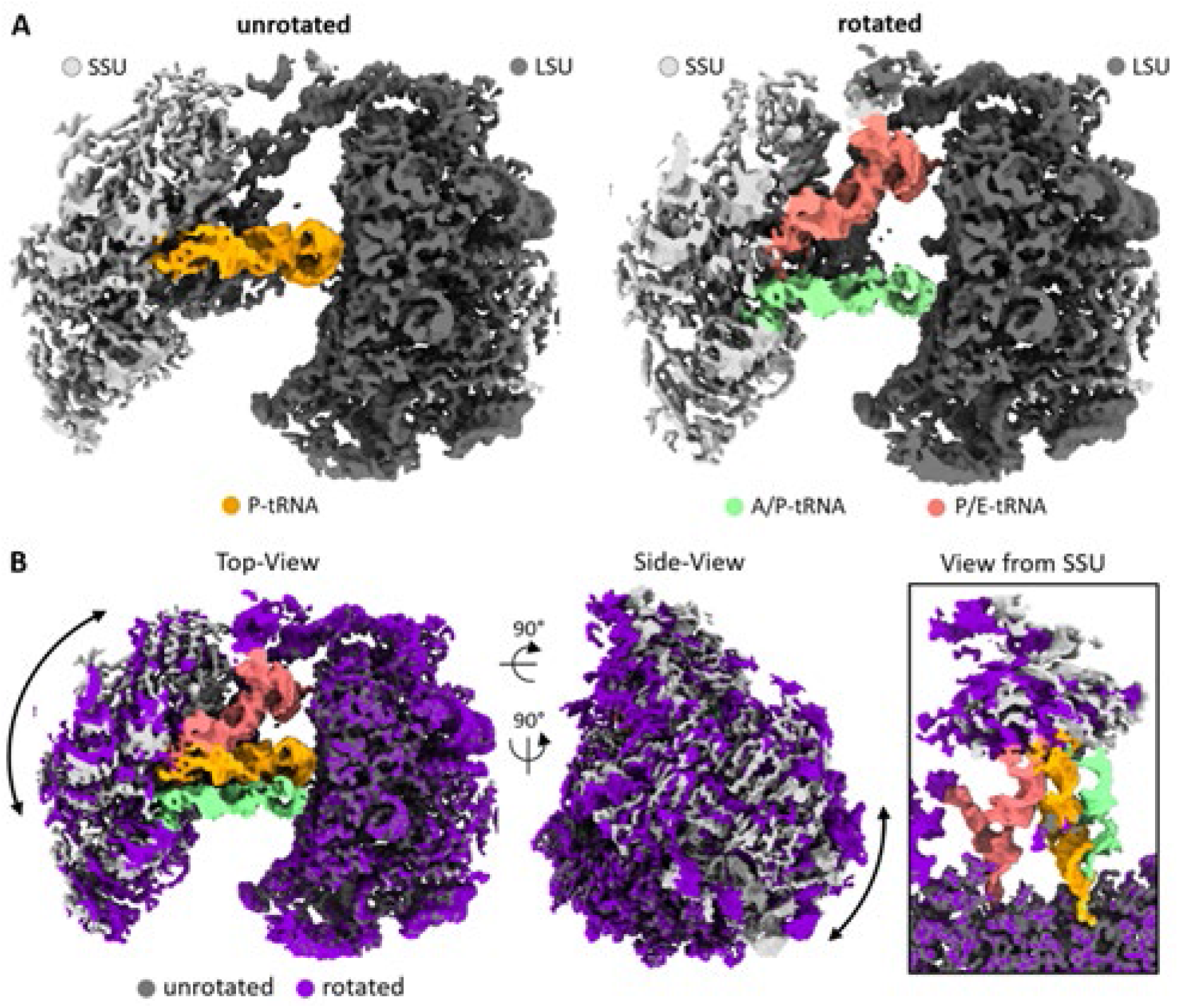
Sucrose-mediated stalling of bZIP11-uORF2 results a rotated and unrotated state of the ribosome. **A**) Cryo-EM maps of the unrotated and rotated state, with the P-tRNA (yellow) and A/P- (green) and P/E-tRNAs (red) highlighted respectively. **B**) Overlap of unrotated and rotated state visualizing the motion on the small subunit (SSU), and close-up view of the tRNA cleft viewed from the SSU.

